# Global diversity, recurrent evolution, and recent selection on amylase structural haplotypes in humans

**DOI:** 10.1101/2024.02.07.579378

**Authors:** Davide Bolognini, Alma Halgren, Runyang Nicolas Lou, Alessandro Raveane, Joana L. Rocha, Andrea Guarracino, Nicole Soranzo, Jason Chin, Erik Garrison, Peter H. Sudmant

## Abstract

The adoption of agriculture, first documented ∼12,000 years ago in the Fertile Crescent, triggered a rapid shift toward starch-rich diets in human populations. Amylase genes facilitate starch digestion and increased salivary amylase copy number has been observed in some modern human populations with high starch intake, though evidence of recent selection is lacking. Here, using 52 long-read diploid assemblies and short read data from ∼5,600 contemporary and ancient humans, we resolve the diversity, evolutionary history, and selective impact of structural variation at the amylase locus. We find that amylase genes have higher copy numbers in populations with agricultural subsistence compared to fishing, hunting, and pastoral groups. We identify 28 distinct amylase structural architectures and demonstrate that nearly identical structures have arisen recurrently on different haplotype backgrounds throughout recent human history. *AMY1* and *AMY2A* genes each exhibit multiple duplications/deletions with mutation rates >10,000-fold the SNP mutation rate, whereas *AMY2B* gene duplications share a single origin. Using a pangenome graph-based approach to infer structural haplotypes across thousands of humans, we identify extensively duplicated haplotypes present at higher frequencies in modern day populations with traditionally agricultural diets. Leveraging 533 ancient human genomes we find that duplication-containing haplotypes (i.e. haplotypes with more *amylase gene* copies than the ancestral haplotype) have increased in frequency more than seven-fold over the last 12,000 years providing evidence for recent selection in West Eurasians. Together, our study highlights the potential impacts of the agricultural revolution on human genomes and the importance of long-read sequencing in identifying signatures of selection at structurally complex loci.

---

Dietary changes have played a major role in human adaptation and evolution impacting phenotypes such as lactase persistence^1,2^ and polyunsaturated fatty acid metabolism^3–5^. One of the most substantial recent changes to the human diet is the shift from hunter-gatherer societies to agricultural-based subsistence. The earliest instance of crop domestication can be traced to the Fertile Crescent of South Western Asia ∼12 thousand years before present (kyr BP) laying the foundation for the Neolithic revolution^6^. Agriculture subsequently spread rapidly westward into Europe by way of Anatolia by ∼8.5 kyr BP and eastward into the Indian subcontinent. However, the transition to agriculture-based subsistence has happened independently several other times throughout human history and today the overwhelming majority of carbohydrates consumed by humans are derived from agriculture.

Plant-based diets are rich in starches which are broken down into simple sugars by *α*-amylase enzymes in mammals. Human genomes contain three different amylase genes located proximally to one another at a single locus: *AMY1,* which is expressed exclusively in salivary glands, and *AMY2A* and *AMY2B,* which are expressed exclusively in the pancreas. It has long been appreciated however, that the amylase locus exhibits extensive structural variation in humans^7,8^ with all three genes exhibiting copy number variation. Indeed, the haplotype represented in the human reference genome GRCh38 contains three tandemly duplicated *AMY1* copies (see methods for details on *amylase* gene naming conventions). Other great apes do not exhibit copy number variation and harbor just a single copy each of the *AMY1, AMY2A,* and *AMY2B* genes^9^. These three amylase genes are the result of duplication events occurring first in the common ancestor of Old World monkeys and apes, and again in the common ancestor of great apes^10^. This ancestral single copy state has also been reported in Neanderthals and Denisovans^11^. *AMY1* copy number correlates with salivary amylase protein levels in humans, and an analysis of seven human populations found increased *AMY1* copy number in groups with high starch diets^12^. While it has been proposed that this gene expansion may have been an adaptive response to the transition from hunter-gatherer to agricultural societies, evidence of recent selection at this locus has been lacking^11,13^. Moreover, subsequent analyses identifying a putative association of *AMY1* copy number and BMI^14^ failed to replicate^15^, highlighting the challenges associated with studying structurally variable loci which are often poorly tagged by nearby single nucleotide polymorphisms (SNPs)^16^. Another major challenge in characterizing selective signatures at structurally complex loci is the difficulty of phasing copy numbers onto haplotypes. Furthermore, while the human reference genome contains a single fully resolved amylase haplotype, the sequence, structure, and diversity of haplotypes on which different copy numbers have emerged are unknown.

## Worldwide distribution of amylase diversity and increased copy number in traditionally agricultural societies

While extensive copy number variation has been documented at the amylase locus in humans^11,14,15,17^, sampling of human diversity worldwide has been incomplete. To explore diversity at this locus we compiled 4,292 diverse high-coverage modern genomes from several sources^18–20^ (see methods for information on all datasets used in this paper) and used read-depth based approaches (see methods, **Fig S1**) to estimate diploid copy number in 147 different human populations (**Figs 1A-C, Extended Data Fig 1, Table S1,** subcontinental groupings as per Mallick et al 2016^20^). Diploid *AMY1* copy number estimates ranged from 2-20 and were highest in populations from Oceanic, East Asian, and South Asian subcontinents. Nevertheless, individuals carrying high *AMY1* copy numbers were present in all continental subgroups. *AMY2A* (0-6 copies) showed the highest average copy number in African populations with deletions more prevalent in non-African populations. *AMY2B* (2-7 copies) exhibited high population stratification with duplications essentially absent from Central Asian/Siberian, East Asian, and Oceanic populations. We also assessed three high coverage Neanderthals and a single Denisovan individual, confirming all to have the ancestral copy number state (**Extended Data Fig 1**). Thus, copy number variation across all three amylase genes is likely human specific.

**Figure 1-.**
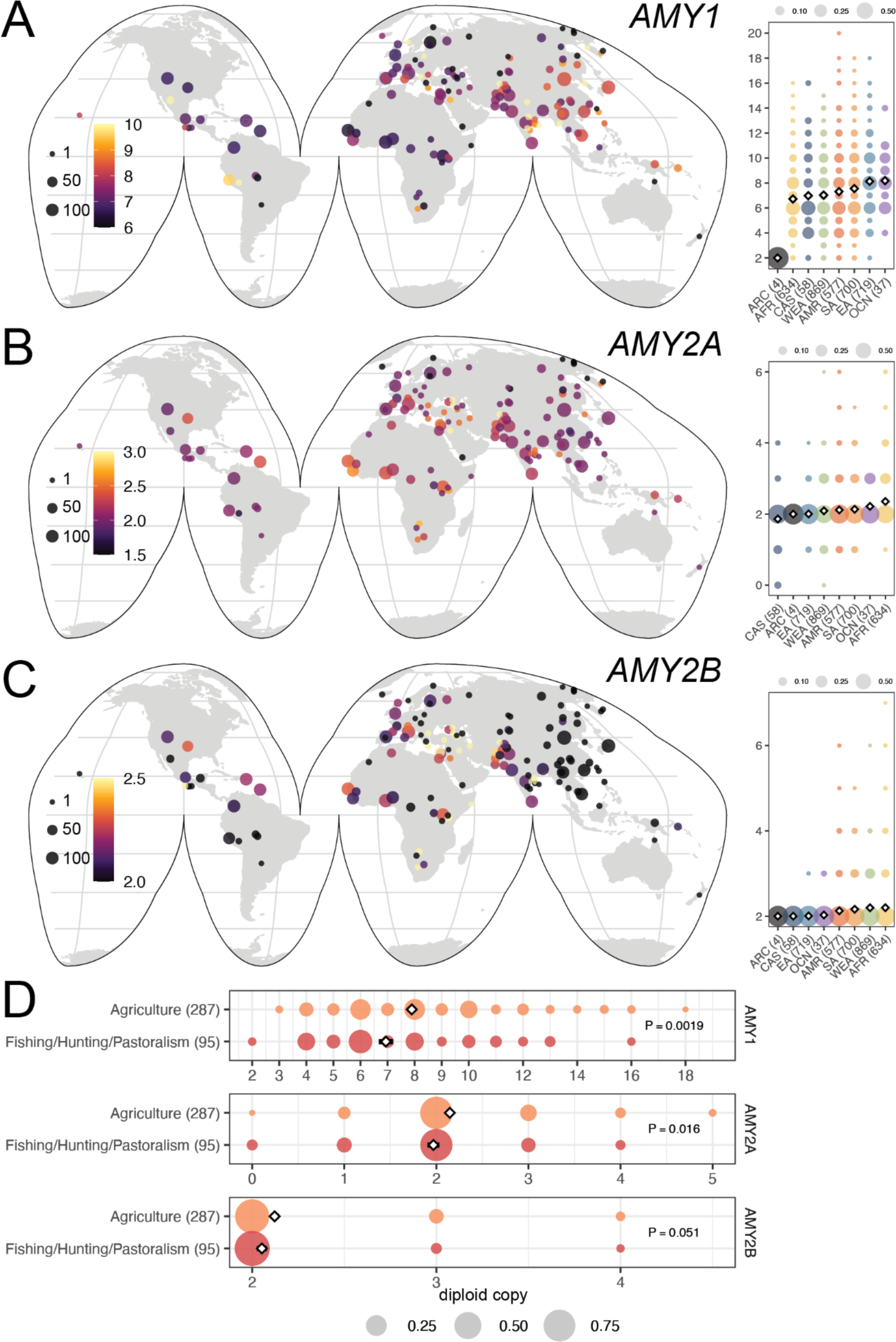
Worldwide amylase copy number diversity. **A-C)** World maps indicating average *AMY1, AMY2A*, and *AMY2B* copy number in 162 different human populations. Point size indicates population sample sizes (ranging from 1-134) and color indicated mean copy number. Inset right are distribution of copy numbers (Y-axis) in continental populations (X-axis): archaic (ARC), African (AFR), Central Asia Siberia (CAS), West Eurasia (WEA), Americas (AMR), South Asia (SA), East Asia (EA), and Oceania (OCN). White diamonds indicate mean, dot sizes indicate proportion with copy number genotype. Copy number distributions across individual populations are displayed in **Extended Data Figure 1. D)** Copy number distributions of *AMY1, AMY2A*, and *AMY2B* in 33 modern human populations with traditionally agricultural subsistence compared to fishing, hunting, and pastoralism-based diets.

While *AMY1* copy number has been shown to exhibit a strong positive correlation with salivary protein levels^12,21^, the relationship between pancreatic amylase gene expression and copy number has not been assessed. Analyzing GTEx^22^ data we confirmed *AMY2A* and *AMY2B* expression was confined to the pancreas. We then genotyped diploid copy numbers in 305 samples for which expression data was available alongside high coverage genome sequencing. Both *AMY2A* (0-5 copies) and *AMY2B* (2-5 copies) copy numbers were significantly and positively correlated with gene expression levels (P=4.4×10^-5^ and P=6.5×10^-4^ respectively, linear model, **Fig S2**).

The strongest evidence of potential selection at the amylase locus comes from comparisons of seven modern day populations with high versus low starch intake^12^. We identified 382 individuals from 33 different populations with traditionally agricultural-, hunter-gatherer-, fishing-, or pastoralism-based diets in our dataset (**Table S2**). The copy number of all three amylase genes was higher in populations with agricultural subsistence compared to those from fishing, hunting, and pastoral groups, though was only strongly significant for *AMY1* (**Fig 1D, S3**, P=0.0019, 0.016, and 0.051 for *AMY1*, *AMY2A*, and *AMY2B* respectively, t-test). These results thus corroborate previous work and demonstrate that pancreatic amylase gene duplications are also more common in populations with starch-rich diets.

## Pangenome-based identification of 28 distinct structural haplotypes underlying extensive amylase copy number variation

The amylase structural haplotype present in the human reference genome (GRCh38) spans ∼200kb and consists of several long, nearly identical segmental duplications. While the approximate structures of several other haplotypes have been inferred through in-situ hybridization and optical mapping, these lack sequence and structural resolution^7,8,12,15^. Nevertheless, the variegated relationship between different amylase gene copy numbers (**Fig 2A**) indicates the existence of a wide range of structures.

**Figure 2-.**
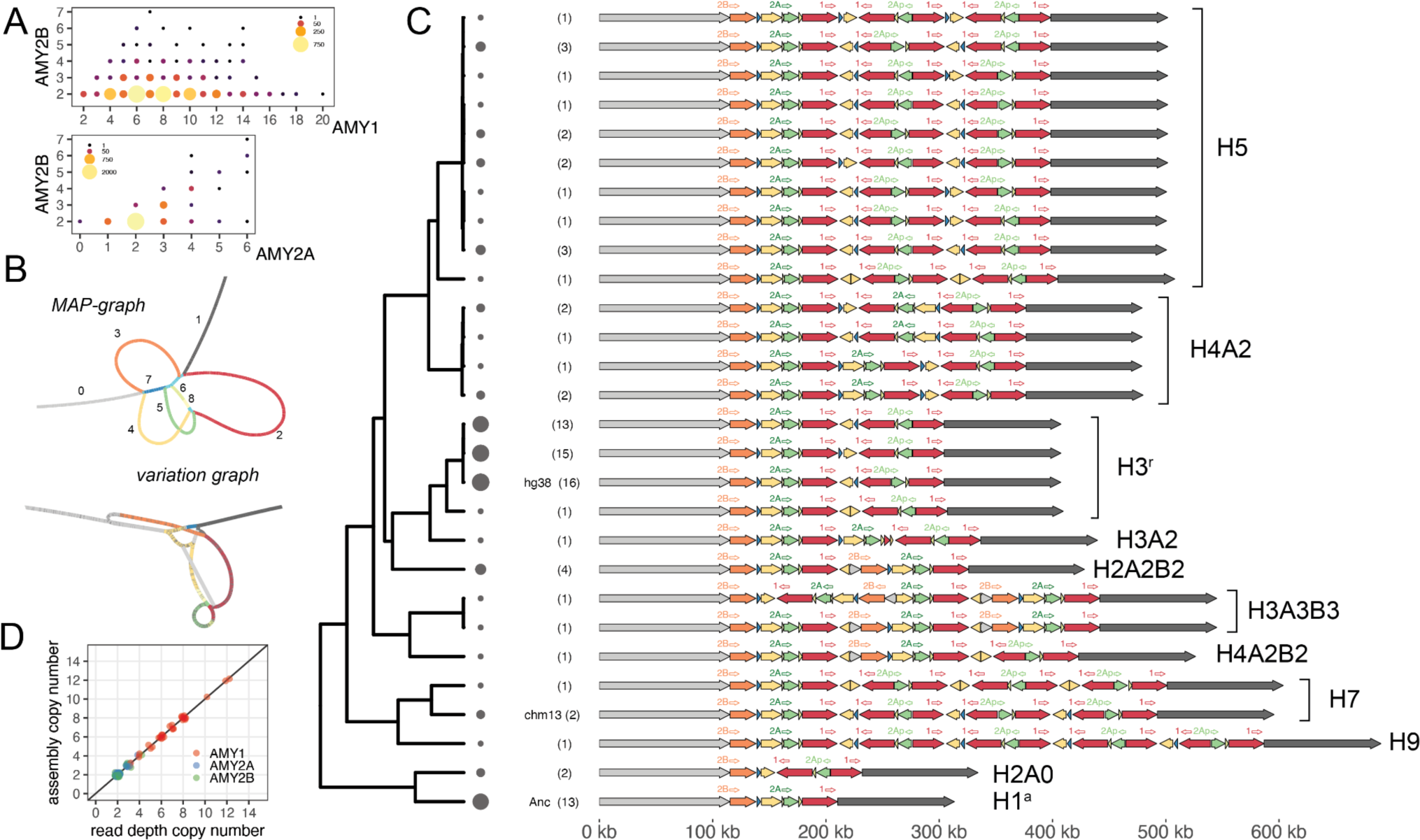
Pangenome-based identification of amylase structural haplotype diversity. **A)** The relationship between *AMY1, AMY2A*, and *AMY2B* copy number. Size and color indicate number of individuals with copy number genotype pair. **B)** Hierarchical minimizer anchored pangenome graph (MAP-graph) and variation graph architectures. Colors and numbers in MAP-graph correspond to principal bundles shown in C. **C)** 28 distinct amylase structural haplotypes identified in 94 haplotypes. Filled arrows indicate principal bundles representing homology relationships while labeled open arrows (above) indicate genes (1 indicates *AMY1*, 2A indicates *AMY2A*, etc.). Numbers in parentheses and circle sizes indicate the number of haplotypes identified with a specific structure. Haplotypes are ordered by their relationship in tree (left) which is generated from the jaccard distance between haplotypes from the variation graph. *Consensus structures*, referring to clusters of similar structures, are indicated to the right. *Consensus structures* names are formatted “H*x*A*y*B*z*”, where *x* corresponds to the copy number of *AMY1*, *y* to the number of *AMY2A*, and *z* to the number of *AMY2B*. “A*y*” and “B*z*” are only included in the name when y or z does not equal to 1. **D)** The relationship between read-depth based copy number and assembly-based copy numbers for amylase genes for 35 individuals (70 haplotypes) in which both haplotypes were assembled across the amylase region.

To characterize the structural diversity of the amylase locus, we first constructed a minimizer anchored pangenome graph (MAP-graph)^23^ from 94 amylase haplotypes derived from 54 long-read, haplotype resolved genome assemblies recently sequenced by the Human Pangenome Reference Consortium (HPRC)^24^ alongside GRCh38 and the newly sequenced T2T-CHM13 reference^25^ (**Fig 2B**, see methods). The MAP-graph captures large-scale sequence structures with vertices representing sets of orthologous or paralogous sequences; thus, input haplotypes can be represented as paths through the graph. We next performed a “principal bundle decomposition” of the graph, which identifies stretches of sequence that are repeatedly traversed by individual haplotypes (colored loops in **Fig 2B**). These principal bundles represent the individual repeat units of the locus. We identified 8 principal bundles in the amylase graph corresponding to: the unique sequences on either side of the structurally complex region containing amylase gene duplications (bundles 0 and 1), the repeat units spanning each of the three amylase genes and the *AMY2Ap* pseudogene (bundles 2, 3, and 5), as well as several other short repeat units (**Fig 2C**). For 35 individuals in which both haplotypes were incorporated into the graph, short read-based diploid genotypes were identical to the sum of the haplotype copy numbers, highlighting the concordance of both short-read genotypes and long-read haplotype assemblies (**Fig 2D**, methods).

Together we identified 28 unique structural haplotypes at the amylase locus (**Fig 2C, Table S3**), of which only 2 had been previously fully sequenced and characterized (the chimpanzee and human reference genome haplotypes). The structurally variable region of the locus (hereafter SVR) spans across all of the amylase genes and ranges in size from ∼95kb to ∼471kb, in all cases beginning with a copy of *AMY2B* and ending with a copy of *AMY1.* To better understand the relationships between these structural haplotypes, we constructed a pangenome variation graph using the PanGenome Graph Builder (PGGB) (**Fig 2B**)^26^. In contrast to the MAP-graph, this graph enables base-level comparisons between haplotypes. Using this graph we computed a distance matrix between all structural haplotypes and built a neighbor-joining tree from these relationships (methods, **Fig 2C**). This tree highlights 11 different clusters of structures each defined by a unique copy number combination of amylase genes (**Fig 2C** right, cluster names correspond to the copy number of AMY1/2A/2B genes, see figure legend for details). Distinct structural haplotypes within each cluster differed largely in the orientation of repeats, or only slightly in their composition. Within each cluster, we assigned one representative structural haplotype as the *“consensus”*. Several of these *consensus* structural haplotypes correspond to approximate architectures which have been previously hypothesized^15^, however 3 of them are described here for the first time (H9, H3A2, and H3A3B3). Among these consensus structures, *AMY1* ranged from 1 to 9 copies with copy 6 and copy 8 states unobserved, *AMY2A* ranged from 0 to 3 copies, *AMY2Ap* ranged from 0 to 4 copies, and *AMY2B* ranged from 1 to 3 copies. We additionally assessed these haplotypes for mutations that might significantly disrupt the function of any of the amylase genes. We identified a single base substitution that introduced a premature stop codon in *AMY1* shared between two haplotypes with high *AMY1* copy number, as well as several missense mutations in all three amylase genes of varying predicted impact (**Table S4**). These mutations were generally found at low frequencies. Because of the low frequency (∼2%) and single origin of the loss-of-function mutation, we do not explicitly account for it in downstream analyses. Together these results reveal the wide ranging and nested-nature of diversity at the amylase locus: different haplotypes can harbor vastly different copy numbers of each of the three genes, and haplotypes with identical gene copy numbers exist in a wide array of forms.

## Time-calibrated inference of haplotype evolutionary histories reveals rapid and recurrent evolution of amylase structures

To discern the evolutionary origins of the vast diversity of structures observed, we sought to explore the SNP haplotypes on which they emerged. We leveraged unique sequences (bundles 0 and 1) flanking the SVR in which SNPs can be accurately genotyped. We first quantified linkage disequilibrium (LD) around the amylase locus in 3,395 diverse human samples (see methods). To our surprise LD was extremely high between SNPs spanning the SVR (∼190-370 kb apart in GRCh38, **Fig 3A, Extended Data Figs 2A-B**). Notably, LD was 7 to 20-fold higher when compared to similarly spaced pairs of SNPs across the remainder of chromosome 1 in all major continental populations (**Fig 3B**). Trio-based recombination rate estimates also indicate reduced recombination rates across the SVR (**Fig 3A** bottom panel)^27^. We hypothesize that these exceptionally high levels of LD arise from the suppression of crossovers between homologs containing distinct structural architectures with vastly different lengths during meiosis^28^.

**Figure 3 -.**
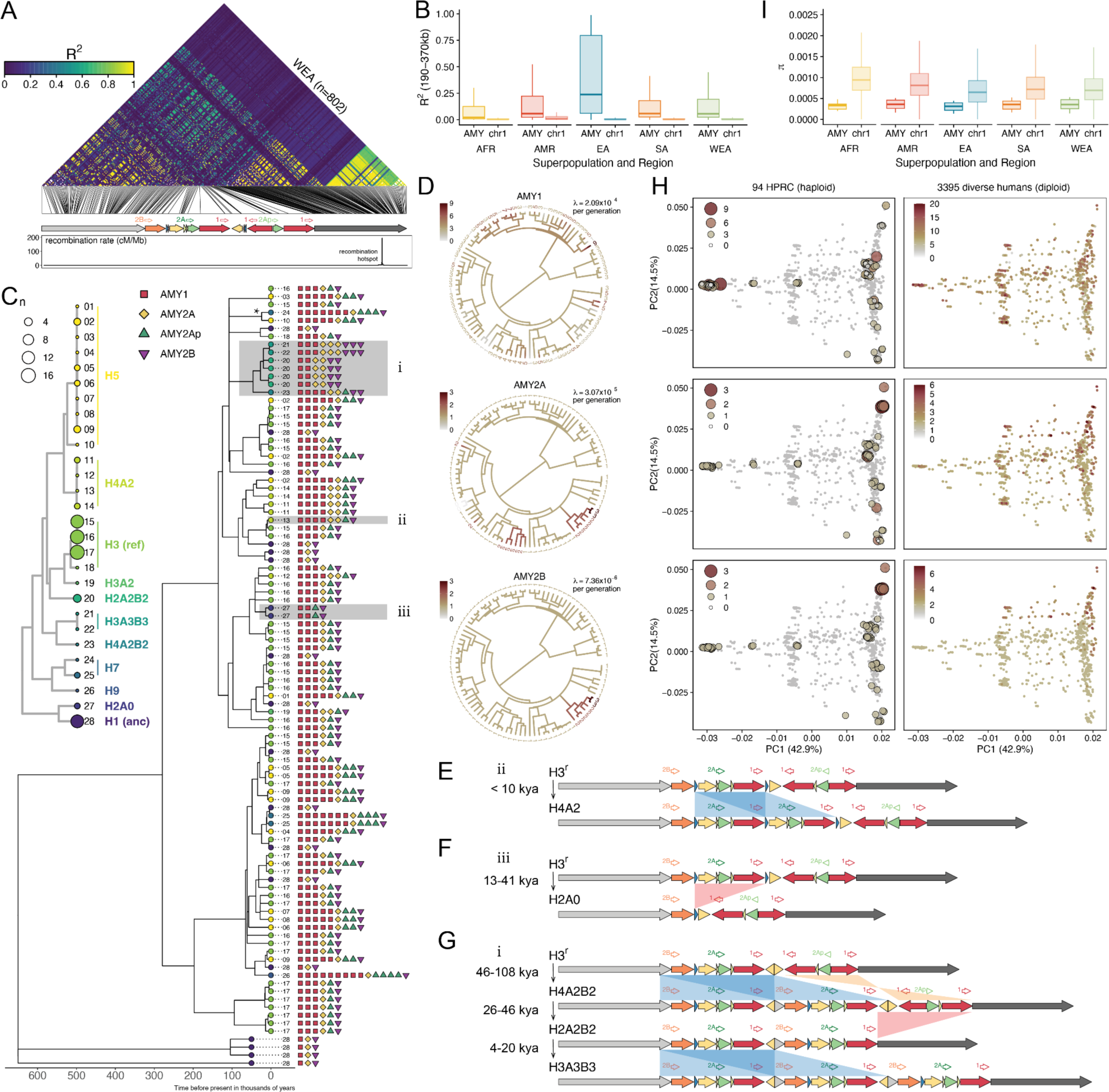
Evolutionary history of amylase structural haplotypes. **A)** Heat map of linkage disequilibrium (LD) for SNPs across a ∼406 kb region spanning unique sequences on either side of the structurally variable region of amylase (SVR) for 802 West Eurasians (WEA) (see **Extended Data Fig. 2A** for all populations). Schematic of GRCh38 structure and recombination rate are shown below. Note that regions outside of the annotated recombination hotspot have recombination rates lower than 0.2 cM/Mb. **B)** Boxplots comparing LD between pairs of SNPs on either side of the SVR (i.e. 190 kb - 370 kb apart) to identically spaced SNPs across chromosome 1 for major human populations with more than 100 samples (see **Extended Data Fig. 2B** for LD decay over genomic distances). **C)** A time-calibrated coalescent tree from the distal non-duplicated region flanking the SVR (leftmost gray arrow in A) across 94 assembled haplotypes (tree from the proximal region in **Extended Data Fig. 3**). The number next to each tip corresponds to the structural haplotype that the sequence is physically linked to and the color of the circle at each tip corresponds to its consensus haplotype structure (see inset structure tree). The copy numbers of each amylase gene and pseudogene are also shown next to the tips of the tree. Asterisk (*) indicates the single, recent origin of the premature stop codon in *AMY1*. **D)** Ancestral state reconstruction and mutation rate estimates for amylase gene copy number (archaic outgroups excluded). Branch color corresponds to copy number. **E-G)** Illustrations of the most recent AMY2A gene duplication, the complete loss of AMY2A gene, and the sequential and joint duplication of AMY2A and AMY2B genes (shaded in gray in C). **H)** A PCA from 94 haplotype assemblies and 3,395 diverse diploid human genomes from the distal non-duplicated region flanking the SVR (PCA from the proximal region in **Extended Data Fig. 3**). In the left column diploid genomes are shown in gray while assembled haplotypes are colored and sized by their haploid amylase copy number. In the right column assembled haplotypes are hidden and diploid genomes are colored by their diploid copy number. **I)** Boxplots comparing *π* calculated in 20 kbp sliding windows across the distal non-duplicated region adjacent to the SVR for major continental human populations with more than 100 individuals.

The high LD across the amylase locus implies that the evolutionary history of the flanking regions are a good proxy for the history of the linked complex structures of the SVR. As such, we constructed a maximum likelihood coalescent tree from these blocks using three Neanderthal haplotypes and a Denisovan haplotype (all containing the ancestral structural haplotype) as outgroups (**Fig 3C**, **Extended Data Fig 3A, S4**, methods). Time calibration of the tree was performed using an estimated 650 kyr BP human-Neanderthal split time^29^. Annotating this coalescent tree with the different amylase structural architectures strikingly revealed that most haplotype structures have experienced repeated evolution, where similar and even identical structures have arisen recurrently on different haplotype backgrounds. Only a handful of structural haplotypes are exceptions to this recurrence, including those harboring *AMY2B* gene duplications which stem from a single originating haplotype.

Our time calibrated tree further enabled us to perform an ancestral state reconstruction for each of the amylase gene copy numbers to quantify the number of times each gene has undergone duplication or deletion (**Fig 3D, Extended Data Fig 3B, S5**). We found that all amylase structural haplotypes in modern humans are descended from an H3^r^ haplotype ∼279 kyr BP. This suggests that the initial duplication event, from the ancestral H1^a^ haplotype to H3^r^, significantly predates the out-of-Africa expansion (i.e. >279 kyr BP). We identified 26 unique *AMY1* gene duplications and 24 deletions since then corresponding to a per generation mutation rate (λ) of 2.09×10^-4^. Although these estimates may be impacted by rare recombination events or additional unsampled duplications/deletions, their magnitude highlights the exceptional turnover of this locus in recent evolution with *AMY1* gene copy number changes occurring at a rate ∼10,000-fold the genome-wide average SNP mutation rate^30^. *AMY2A* exhibited substantially fewer mutational events, undergoing 6 duplications and 2 deletions (λ=3.07×10^-5^) with the most recent *AMY2A* duplication occurring within the last 9.4 kyr BP (**Figs 3D, E**). While duplications of *AMY2A* have occurred several times, we identified a single origin of the complete loss of the *AMY2A* gene in our tree, which occurred 13.5-40.7 kyr BP and resulted in the H2A0 haplotype (**Figs 3D, F**). Only 2 *AMY2B* duplications were identified (λ=7.36×10^-6^), occurring sequentially on a single haplotype and thus allowing us to resolve the stepwise process of their formation (**Figs 3D, G**). We estimate the first duplication event occurred 46-107.8 kyr BP, followed by a deletion 26.9-46 kyr BP, and finally by a second duplication event 4.1-19.5 kyr BP (**Fig 3G**).

While our collection of 94 assembled haplotypes spanning the complex SVR provides the most complete picture of amylase evolution to date, it still represents just a small fraction of worldwide genetic variation. To characterize the evolution of amylase haplotypes more broadly, we performed a PCA combining the fully assembled haplotypes with 3,395 diverse human genomes using the flanking regions of the SVR (**Fig 3H, Extended Data Fig 3C, S6, S7**). We annotated individuals in the PCA with haploid/diploid *AMY1/2A/2B* copy numbers respectively. As expected, clusters of diploid individuals with high copy number (**Fig 3H** right panels) tended to colocalize with assembled haplotypes containing duplications (**Fig 3H** left panels). Exceptions to this indicate heterozygotes (with placements in between two haplotypes) or additional duplication/deletion events. This method identified several additional *AMY1* and *AMY2A* duplication events worldwide, as expected given their high mutation rate, and support for additional haplotypes with complete *AMY2A* deletions (**Figs 3H, S6**). However, we find no evidence of additional *AMY2B* gene duplications, supporting the single origin of these haplotypes.

## Reconstruction of complex amylase structures from short read data uncovers worldwide diversity, stratification, and haplotypes associated with agriculture

Our analyses of SNP diversity at regions flanking the amylase SVR also revealed a substantial reduction in diversity compared to the chromosome-wide average (quantified by *π*, 2-3 fold lower, **Fig 3I**). To further investigate if this signature was indicative of a selective sweep we ran several genome-wide selection scans (iHS^31^, nSL^32^, H12 and H2/H1^33^, composite likelihood *saltiLASSI* statistic^34^, XP-nSL^35^, **Table S5, Figs S8-S19**). We found that the nSL and H2/H1 statistics tended to be higher at regions flanking the amylase SVR in specific populations (WEA, CAS, and modern populations with traditionally agricultural diets), consistent with a soft or incomplete sweep. However, these results fell below the 99.95% threshold of the genome-wide empirical distribution^31^, though this could be a consequence of the limitations of SNP-based methods in detecting selection at rapidly-evolving, structurally complex loci, where identical structures repeatedly emerge on distinct haplotype backgrounds.

Instead of relying on neighboring SNPs as a proxy for amylase structural variants, we developed an approach to directly identify the structural haplotype pairs present in short-read sequenced individuals. Briefly, this approach, which we term *‘haplotype deconvolution’*, consists of mapping a short read-sequenced genome to the pangenome variation graph (**Fig 4A**) and quantifying read depth over each node in the graph (n=6,640 nodes in the amylase graph). This vector of read depths is then compared with a set of precomputed vectors generated by threading all pairs of 94 long-read assembled haplotypes (i.e., all possible genotypes) over the same graph. Finally, we infer the structural genotype of the short read genome to be the pair of long-read assembled haplotypes whose vector representation most closely matches to the short-read vector (**Fig 4B**, see methods). We assessed the accuracy of this approach using three orthogonal methods. First, we compared haplotype deconvolutions in 35 individuals for which both short-read data and haplotype-resolved assemblies were available. Short read-based haplotype deconconvolutions exactly matched the long read assembly haplotypes 100% of the time (70/70 haplotypes). Second, we used 602 diverse short-read sequenced trios and estimated the accuracy of haplotype inference to be ∼94% from Mendelian inheritance patterns (see methods) and 95%-97% concordant with previous inheritance-based determinations of haplotypes in 44 families^15^. Finally, we compared our previously estimated reference genome-based copy number genotypes to those predicted from *haplotype deconvolutions* across 4,292 diverse individuals. These genotypes exhibited 95-99% concordance across different amylase genes (95%, 97%, and 99% for *AMY1, AMY2A*, and *AMY2B* respectively). Cases in which the two estimates differed were generally rare high-copy genotypes for which representative haplotype assemblies have not yet been observed and integrated into the graph (**Fig S20**). Thus, we determine that our *haplotype deconvolution* method is robust and ∼95% accurate, and limited primarily by the completeness of the reference pangenome.

**Figure 4 -.**
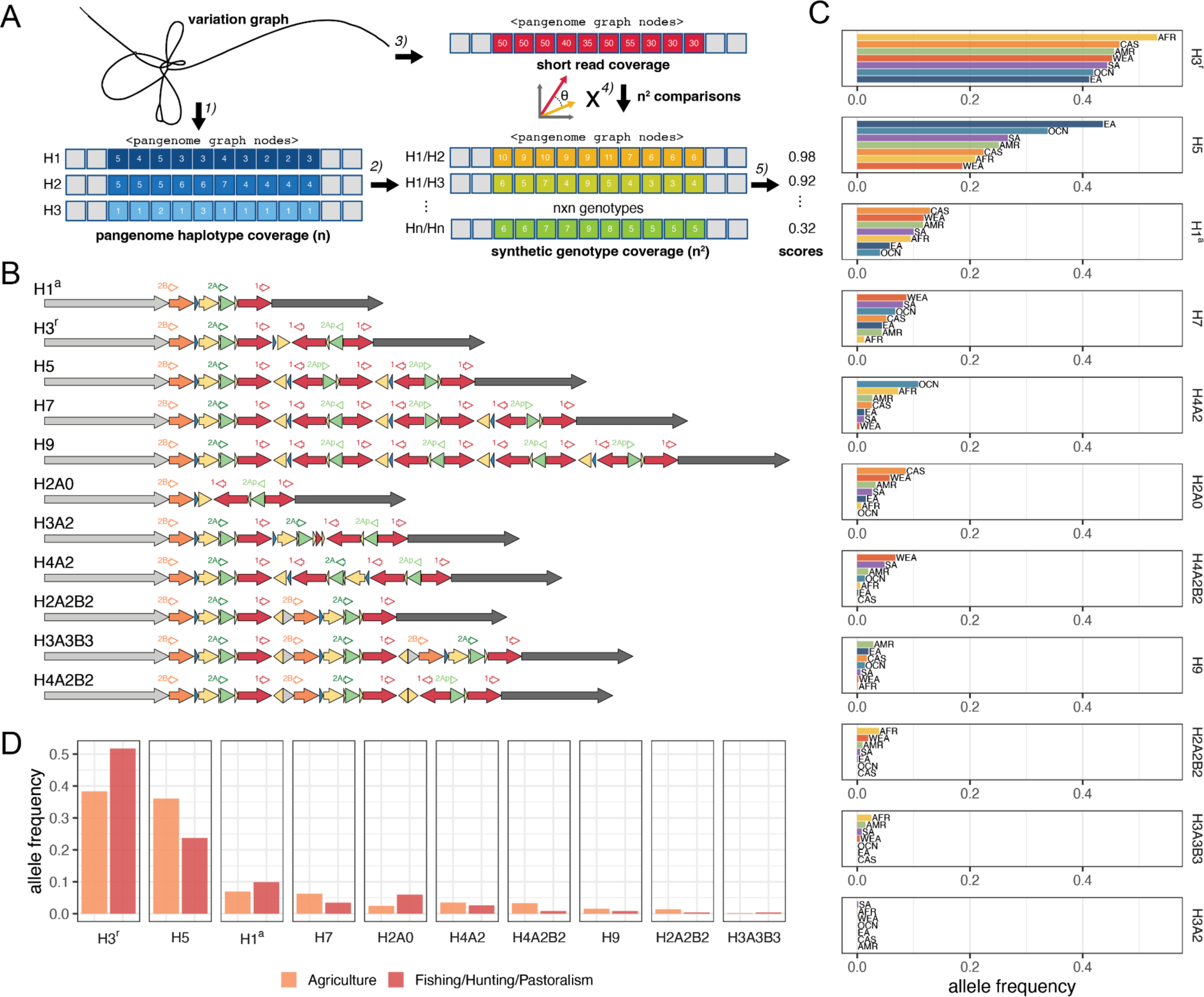
Inference of complex structural haplotypes from short-read data. **A)** A schematic of the *haplotype deconvolution* approach to infer the pair of structural haplotypes present in a short-read sequenced individual. 1) A set of assembled haplotypes are mapped to a variation graph and coverage vectors are quantified over all nodes of the graph. 2) Synthetic genotype vectors are constructed from summing all pairs of haplotype vectors. 3) A short-read genome is mapped to the variation graph and read depth is quantified over all nodes in the graph. 4) The short-read coverage vector is compared to all synthetic genotype vectors and scored (5) to identify the most likely haplotype pair present in the short-read sequenced individual. **B)** Consensus haplotype structures. **C)** Structural haplotype frequencies across continental populations in 3,594 diverse humans (7188 haplotypes). **D)** Haplotype (allele) frequencies in individuals with traditionally agricultural subsistence compared to fishing, hunting, and pastoralism based diets.

We used *haplotype deconvolution* to estimate worldwide allele frequencies and continental subpopulation allele frequencies for amylase consensus structures across 7,188 haplotypes (**Figs 4B, C, Tables S6, S7**). The reference haplotype, H3^r^, was the most common globally however several haplotypes exhibited strong population stratification. The H5 haplotype is the most frequent haplotype in East Asian populations whereas the ancestral haplotype H1^a^ was underrepresented in East Asian and Oceanic populations. The high copy H9 haplotype was largely absent from African, West Eurasian, and South Asian populations, while ranging from 1-3% in populations from the Americas, East Asia, and Central Asia and Siberia. Haplotypes with *AMY2B* duplications (i.e. H2A2B2, H3A3B3, and H4A2B2) were essentially absent from East and Central Asia, explaining our previous observation of the lack of *AMY2B* duplication genotypes in these global populations (**Fig 1C**) and consistent with their single origin.

We next compared the relative haplotype frequencies among modern human populations with traditionally agricultural-, hunter-gatherer-, fishing-, or pastoralism-based diets (**Fig 4D**). Agricultural populations differed significantly from non-Agricultural populations (p=0.011, chi squared test) and were enriched for haplotypes with higher *AMY1* copy number, including the H5, H7, and H9 haplotypes, as well as for haplotypes with higher *AMY2A* and *AMY2B* copy number (H4A2B2, H2A2B2). In contrast, fishing, hunting, and pastoralism-based populations were enriched for the reference H3^r^, deletion H2A0, and ancestral H1^a^ haplotypes. These results demonstrate that haplotypes with increased amylase gene copy number are enriched in modern day populations with traditionally agricultural diets.

## Ancient genomes reveal recent selection at the amylase locus in West Eurasian populations

The development of agriculture ∼12,000 years ago in the Fertile Crescent catalyzed a rapid shift in the diets and lifestyles of West Eurasian populations. Most of the ancient genome sampling to date has been performed in Europe, allowing us to deeply explore the evolution of the amylase locus in these populations following the adoption of agriculture. To uncover how the genetic diversity of the amylase locus was shaped over this time period we collated 533 recently generated ancient genomes from West Eurasia^36,37^, ___which span in age from ∼12,000 to ∼250 BP (**Figs 5A, S21, Table S8**). We estimated amylase gene copy numbers from these ancient individuals and compared these with copy numbers in modern Europeans (**Figs 5B, S22, Table S1**). Overall, copy numbers of all amylase genes tended to be lower in ancient hunter gatherer populations compared to Bronze Age through present day European populations, although these comparisons are of varying statistical significance due to our limited sample size of some ancient populations (ANOVA followed by Tukey’s test, **Fig 5B, Table S9**). We next assessed how total copy numbers have changed as a function of time for each of the three amylase genes (**Fig 5C**). In all three cases we observed significant increases in total copy number over the last ∼12,000 years (P=1.1×10^-6^, 1.6×10^-6^ and 0.0032 for *AMY1*, *AMY2A*, and *AMY2B* respectively, linear model). The total *AMY1* copy number increased by an average of ∼2.9 copies over this time period while *AMY2A* and *AMY2B* increased by an average of 0.4 and 0.1 copies respectively. These results are suggestive of directional selection at this locus for increased copy number of each of the three amylase genes.

**Figure 5-.**
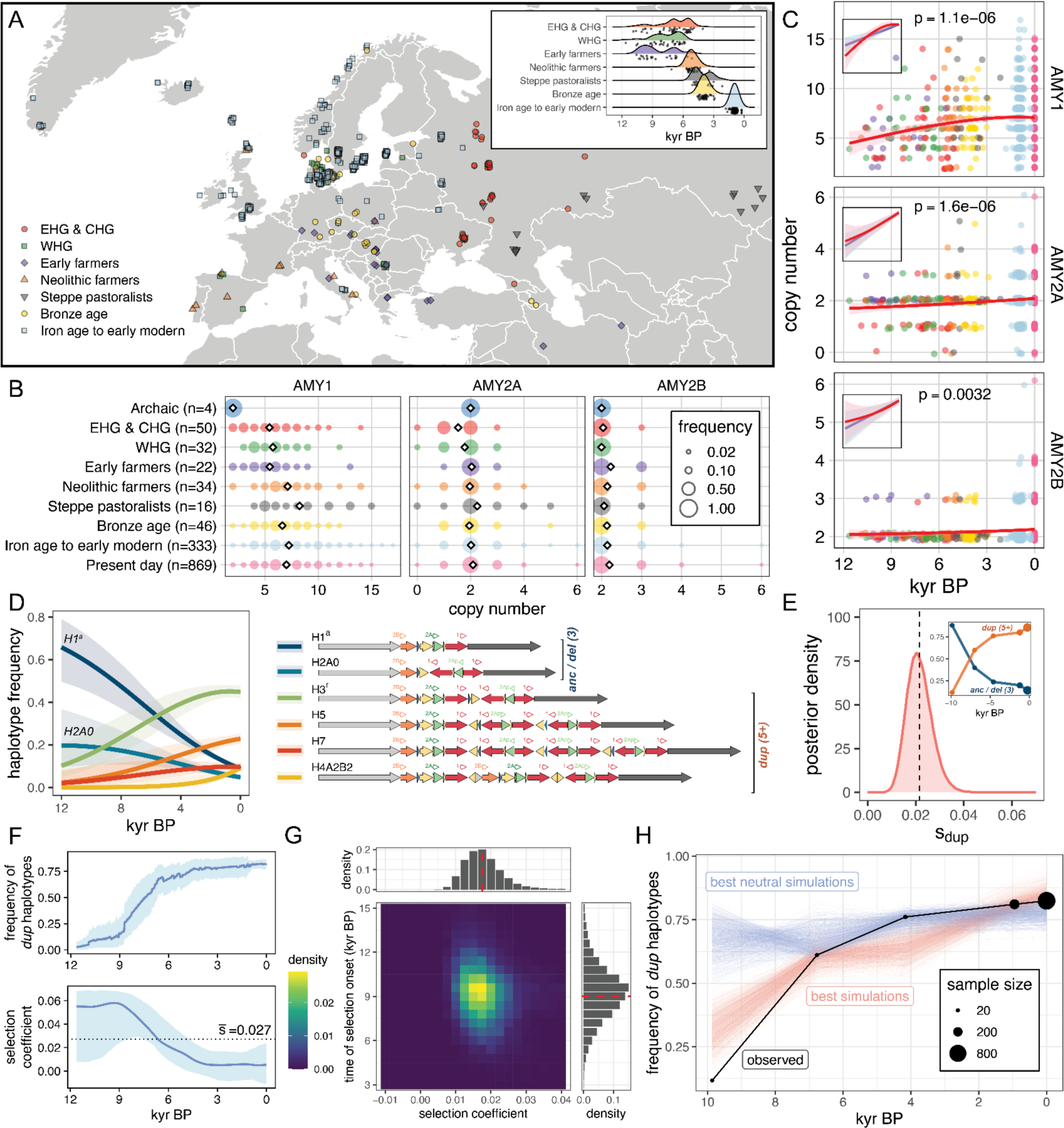
Recent selection at the amylase locus in West Eurasia. **A)** Locations of 533 West Eurasian ancient genomes from which amylase copy numbers were estimated. Inset shows the estimated ages of these samples. **B)** The distribution of *AMY1, AMY2A*, and *AMY2B* copy numbers in ancient and modern populations of West Eurasia. **C)** Copy number genotypes plotted as a function of age overlaid with a smooth generalized additive model fit. Inset shows isolated linear model (blue) and generalized additive model (red) fit to data. P-values from the linear model are shown. **D)** Haplotype trajectories fit by multinomial logistic regression for 6 haplotypes (right) present at >1% frequency in ancient and modern West Eurasians. Structures with the ancestral 3 total amylase copies (*anc/del*) are distinguished from duplication-containing haplotypes with ≥5 amylase genes (*dup*). **E)** Posterior density of the selection coefficient for *dup* haplotypes over the last 12,000 years estimated from ApproxWF (mean 0.022, indicated by dotted line, no estimates ≤ 0 were observed in 1,000,000 MCMC iterations). Inset are binned observations of *dup* versus *anc/del* haplotype frequency trajectories. **F)** Frequency and selection coefficient trajectories for *dup* haplotypes and their 95% credible interval estimated from *bmws*. **G)** Posterior distribution of the selection coefficient and the time of selection onset based on the ABC approach. Red dashed lines mark the median of the distribution. **H)** The observed allele frequency trajectory and the expected allele frequency trajectories from the top 1000 of all simulations and top 1000 of neutral simulations.

We next applied our haplotype deconvolution approach to these ancient genomes to infer how the frequency of amylase structural haplotypes has changed over recent time. Simulations confirmed this method to be highly accurate even on low-coverage ancient genomes (methods, **Fig S23**). We further conservatively selected 288 of the 533 individuals with the highest confidence haplotype assignments (see methods, **Figs S24-S25**, **Table S6**). Six haplotypes were found at appreciable frequencies (>1%) in either modern or ancient West Eurasian populations including the H1^a^ and H2A0 (*AMY2A* deletion) haplotypes, which each contain 3 total functional amylase gene copies, and the H3^r^, H5, H7, and H4A2B2 haplotypes, which contain between 5 and 9 total amylase gene copies (**Figs 5D, S26**). Modeling the frequency trajectories of each of these haplotypes using multinomial logistic regression, we found that the ancestral H1^a^ and the H2A0 haplotypes both decreased significantly in frequency over the last ∼12,000 years, from a combined frequency of ∼0.88 to a modern day frequency of ∼0.14 (**Figs 5D, 5E inset, S25-S27**). In contrast, duplication-containing haplotypes (with 5 or more amylase gene copies in contrast to the ancestral 3 copies - we note that no haplotypes containing 4 copies are observed) increased in frequency commensurately more than 7-fold (from ∼0.12 to ∼0.86) over this time period.

We used three complementary approaches to test whether positive selection could explain the substantial rise in the frequency of duplication-containing haplotypes (see methods for model parameters and assumptions). First, we used a Bayesian approach that assumes a constant population size and selection coefficient (*ApproxWF*^38^). The posterior distribution of the selection coefficient supported positive selection (P<1×10^-6^, empirical p-value) with an average of s_dup_=0.022 (**Fig 5E**). We next employed *bmws*^39^, which allows s_dup_ to vary over time. Selection was found to be the strongest 12-9 kyr BP, with s_dup_ approaching 0.06 (**Fig 5F**). Subsequently, selection has significantly weakened, approaching 0 in recent times (average s_dup_= 0.027, **Fig 5F**). Lastly, we implemented an approximate Bayesian computation (ABC) approach adapted and modified from Kerner *et al.*^40^ to account for the important demographic factors that shape allele frequencies over time (e.g. population structure, admixture events, population growth, see methods). The posterior distribution of s_dup_ is centered around 0.0175 and does not overlap 0 while the time of the selection onset is estimated to be around 9 kyr BP (**Figs 5G, S28**). In addition, none of the neutral simulations conducted (i.e. with s_dup_=0) exhibits higher allele frequency increases than observed in the data (**Figs 5H, S29)**. Taken together, these results are consistent with positive selection for duplication-containing haplotypes at the amylase locus following the adoption and spread of agriculture in West Eurasia.

## Discussion

The domestication of crops and subsequent rise of farming radically reshaped human social structures, lifestyles, and diets. Several evolutionary signatures of this transition have been identified in ancient and modern West Eurasian genomes^37,41,42^. However, while it has been hypothesized that the amylase locus has similarly undergone selection due to this transition^12^, footprints of recent positive selection have not been detected to date^11,13^. Here, taking advantage of long read assemblies, we characterize the complex haplotype structures at the amylase locus to the highest resolution to date, illuminating structural and sequence complexity intractable to short read sequencing (e.g. **Fig S30**). Furthermore, these long read haplotypes for the first time provide information about flanking SNPs linked to these complex structures. These enable us to build coalescent trees revealing the rapid and repeated duplication and deletion events at this locus in recent human history. In particular, we find that the majority of these events occurred within the last 50KY and thus would only be tagged by rare variants in the flanking region. Thus, the extensive homoplasy and high mutation rate at this region make flanking SNPs poor tags in classical tests for selective sweeps^43,44^, potentially explaining the failure of previous efforts aimed at detecting selection at this locus. Finally, we leverage long read assemblies to improve the utility of existing short read data by constructing pangenome graphs of the amylase locus which we use to infer the haplotype structure in short-read sequenced individuals. This graph-based approach, termed “haplotype deconvolution,” unlocks a new era where regions previously inaccessible to short reads can now be revisited in both modern and ancient datasets.

Using our haplotype deconvolution approach we were able to confidently reconstruct the haplotype structures of 288 ancient samples at the amylase locus. We find that haplotypes carrying duplicated copies of amylase genes have increased in frequency seven-fold in the last 12,000 years. We note that our analyses are limited by the relatively low sample sizes and uneven sampling of high quality ancient genomes in West Eurasia suitable for haplotype assignment. The several approaches we used to test for selection are also dependent on various model assumptions and genotyping accuracy. Nevertheless, we present multiple lines of evidence (**Figs 1D, 4D, 5C-H**) that consistently support recent selection in West Eurasians at the amylase locus potentially linked to the adoption of agriculture.

One of the best studied examples of human adaptation to diet is the evolution of lactase persistence^1,2^ (though see^45,46^ regarding potential complexities underlying selection at this locus). Intriguingly, our estimates of s_dup_ are comparable in magnitude to estimates of s at the *MCM6/LCT* locus reported in many studies^39,40,45,47^. However, increased *AMY1* copy numbers have also been associated with deleterious oral health outcomes^48^ (i.e. cavities), highlighting a potential evolutionary tradeoff which might result in distinct selection dynamics in contrast to other diet associated loci like *LCT*. The repeated mutation and homoplasy found at the amylase locus adds further evolutionary complexity, in contrast to loci driven by point mutations. We find the mutation rate of amylase gene duplications/deletions to be ∼10,000-fold the average SNP mutation rate, similar to short tandem repeats^49^. This is similar to recently described structural variation mutation rates at ampliconic Y chromosome regions^50^. In both cases the duplication architecture of the locus potentially predisposes to de-novo SV formation through non-allelic homologous recombination (NAHR) between long paralogous sequences on the same chromatid or sister chromatids^51–53^, or non-crossover gene conversion which can yield similar SVs^54^. Thus linkage disequilibrium is maintained across the locus even in the presence of rapid, recurrent structural changes resulting in 28 distinct haplotype structures, many of which have multiple origins. Our analyses contrast duplication-continuing versus non-duplication-containing haplotypes as a simplification given our limited sample sizes. However, the interaction of multiple haplotypes and their distinct evolutionary trajectories remains an exciting direction to explore. More broadly, the selective signature associated with duplication-containing amylase haplotypes illustrates the critical role SVs can play in human evolution. SVs can alter gene dosage, reconfigure the heterochromatic landscape of the genome, and reshape patterns of recombination.

Another interesting parallel between MCM6/LCT and amylase is that the ability to digest milk has arisen independently in different populations^1,2^. Similarly, agriculture has been adopted independently several times throughout human history^6^. Here, in addition to showing evidence of positive selection in West Eurasian populations, we find that haplotypes carrying higher amylase copy numbers are found more commonly in multiple other populations with traditionally agricultural subsistence worldwide. These results suggest that selection for increased amylase copy number may have also happened several times throughout human history, coincident with the several independent adoptions of agriculture. Because ancient samples from regions other than Europe are scarce, we were not able to infer potential selection associated with other agricultural adoptions. More extensive sampling of diverse ancient genomes and modern long-read assemblies are needed to further test this hypothesis. Remarkably, the expansion of amylase genes accompanying transitions to starch-rich diets appears to have also occurred independently across several different commensal species including dogs, pigs, rats, and mice, highlighting the repeated evolution of this locus across taxa^9,55^ and the far reaching impact of the agricultural revolution on the genetics and evolution of species beyond our own.

## Online content

Supplementary figures can be found in Supplementary Online Materials

## Methods

### Amylase gene naming conventions

The reference genome GRC38 represents an H3 haplotype with three copies of the *AMY1* gene and one copy each of the *AMY2A* and *AMY2B* genes. The three *AMY1* copies are identified with labels *AMY1A, AMY1B*, and *AMY1C* due to HUGO naming convention requirements for all gene copies to have unique names. However, these various copies of *AMY1* genes across different haplotypes are recent duplications that share high sequence similarity, and therefore are referred to simply as *AMY1* genes in this paper and others^11,12,15^. In contrast, AMY2A and AMY2B stem from a much older gene duplication event and are much more diverged than the different copies of AMY1 genes^10^. They share the AMY2 prefix simply because they are both expressed in the pancreas.

### Datasets

Short-read sequencing data were compiled from high-coverage resequencing of 1000 genome samples^19^, the Simons Genome Diversity Panel^20^, and the Human Genome Diversity Panel^18^. Genomes from GTEx^22^ samples were also assessed, but only for gene expression analyses as the ancestry of these samples was not available. In total, we obtained copy number genotype estimates for 5,130 contemporary samples. Among these, 838 are GTex samples, 698 are trios from the 1000 Genomes Project (1KG), and the rest (n=3,594, i.e. 7,188 haplotypes) are unrelated individual samples compiled from 1KG, HGDP, and SGDP. GTEx and 1KG trio samples were excluded from analyses characterizing the global diversity of the amylase locus. We performed haplotype deconvolutions on all unrelated samples as well as trio data (n=4,292 total), but the trios were only used for validation purposes.

Figure S30 shows SV calls from the gnomAD project^56^. Phased SNP calls from 1000 genomes and HGDP samples were compiled from Koenig et al.^57^, which includes all of our 1KG and HGDP samples but only some of the SGDP samples (n=3,395 total). These data were used for the analyses of LD, nucleotide diversity, PCA, and selection scans^57^.

Ancient genome short-read fastq samples were compiled from Allentoft et al.^37^ and Marchi et al.^36^ and were mapped to the human reference genome GRCh38 with BWA (v0.7.17, ‘`bwa mem`‘)^58^. The modern genomes as well as the 14 Marchi et al. genomes are of high coverage and quality, however the Allentoft et al. samples were of varying quality and coverage. The Allenoft et al. dataset included more than 1600 ancient genomes including 317 newly sequenced ancient individuals alongside 1492 previously published genomes. Unfortunately, many published ancient genomes have been filtered to exclude multi-mapped reads leaving large gaps over regions such as the amylase locus. After removing genomes with missing data, 690 samples remained. We carefully analyzed these 690 genomes to determine their quality by quantifying the standard deviation of genome-wide copy number (after removing the top and bottom fifth percentiles of copy number to exclude outliers). We chose a standard deviation cutoff of 0.49 based on a visual inspection of the copy number data and selected 519 samples (∼75% of 690) with sufficient read depth for copy number genotyping. Ancient samples were assigned to one of eight major ancient populations in West Eurasia based on their genetic ancestry, location, and age obtained from their original publications^3637^ (**Figs 5A, S21, Table S8**). These populations include: Eastern hunter-gatherer (EHG), Caucasian hunter-gatherer (CHG), Western hunter-gatherer (WHG), Early farmer (samples with primarily Anatolian farmer ancestry), Neolithic farmer (samples with mixed Anatolian farmer and WHG ancestry), Steppe pastoralist (samples with mixed EHG and CHG ancestry), Bronze age (samples with mixed Neolithic farmer and Steppe ancestry), and Iron age to early modern. Lastly, four archaic genomes were assessed including three high coverage Neanderthal Genomes and the high-coverage Denisova genome^29,59–61^.

Long-read haplotype assemblies were compiled from the human pangenome reference consortium (HPRC)^24^. Year 1 genome assembly freeze data were compiled along with year 2 test assemblies. Haplotype assemblies were included in our analyses only if they spanned the amylase SVR. Furthermore, in cases where both haplotypes of an individual spanned the SVR, we checked to ensure that the diploid copy number of amylase genes matched with the read-depth based estimate of copy number. We noted that several year 1 assemblies (which were not assembled using ONT ultralong sequencing data) appeared to have been misassembled across the amylase locus as they were either discontiguous across the SVR, or had diploid assembly copy numbers that did not match with short-read predicted copy number. We thus reassembled these genomes incorporating ONT ultralong sequence using the Verkko assembler^62^ constructing improved assemblies for HG00673, HG01106, HG01361, HG01175, HG02148, HG02257. Alongside these HPRC genome assemblies, we included GRCh38 and the newly sequenced T2T-CHM13 reference^25^.

### Determination of subsistence by population

The diets of several populations (see **Table S2**) were determined from the literature from the following sources^12,63–71^. We were able to identify the traditional diets for 33 populations. All other populations were excluded from this analysis.

### Read depth based copy number genotyping

Copy number genotypes were estimated using read depth as described in^16^. Briefly, read depth was quantified from BAMs in 1000bp sliding windows in 200bp steps across the genome. These depths were then normalized to a control region in which no evidence of copy number variation was observed in >4000 individuals. Depth-based “raw” estimates of copy number were then calculated by averaging these estimates over regions of interest. Regions used for genotyping are found in **Table S10**. We note that the AMY2Ap pseudogene is a partial duplication of the AMY2A that excludes the ∼4500bp of the 5’ end of the gene. This region can thus be used to genotype AMY2A copy without “double counting” AMY2Ap gene duplicates. Copy number genotype likelihoods were estimated by fitting modified Gaussian Mixture Model (GMM) to “raw” copy estimates across all individuals with the following parameters: *k -* the number of mixture components, set to be the difference between the highest and lowest integer-value copy numbers observed; *π -* a k-dimensional vector of mixture weights; σ - a single variance term for mixture components; *o -* an offset term by which the means of all mixture components are shifted. The difference between mixture component means was fixed at 1 and the model was fit using expectation maximization (**Fig S1**). The copy number maximizing the likelihood function was used as the estimated copy number for each individual in subsequent analyses. Comparing these maximum likelihood copy number estimates with ddPCR yielded very high concordance with r^2^ = 0.98, 0.99 and 0.96 for *AMY1*, *AMY2A*, and *AMY2B* respectively (**Fig S1**). For comparisons of copy number as a function of sustenance, populations were downsampled to a maximum of 50 individuals. We also employed a linear mixed effects model approach in which all samples were maintained which provided similar results (P=0.013, 0.058, 0.684).

### Analysis of gene expression

Gene expression data from the GTEx project^22^ were downloaded alongside short read data (see above section). Normalized gene expression values for *AMY2A* and *AMY2B* were compared to copy number estimates using linear regression (**Fig S2**).

### Minimizer Anchored Pangenome Graph Construction

Regions overlapping the amylase locus were extracted from genome assemblies in two different ways. First, we constructed a PanGenome Research Tool Kit (PGR-TK) database from HPRC year 1 genome assemblies and used the default parameters of w=80, k=56, r=4, and min-span=64 for building the sequence database index. The GRCh38 chr1:103,655,518-103,664,551 was then used to identify corresponding AMY1/AMY2A/AMY2B regions across these individuals. Additional assemblies were subsequently added to our analysis by using minimap2^72^ to extract the amylase locus from those genome assemblies. The Minimizer Anchored Pangenome Graph and the Principal Bundles were generated using revision v0.4.0 (git commit hash: ed55d6a8). The Python scripts and the parameters used for generating the principal bundle decomposition can be found in the associated GitHub Repository. The position of genes along haplotypes was determined by mapping gene modes to haplotypes using minimap2^72^.

### Analysis of mutations at amylase genes

To identify mutations in amylase genes from long-read assemblies and evaluate their functional impact, we first aligned all amylase gene sequences to *AMY1A*, *AMY2A*, and *AMY2B* sequences on GRCh38 using minimap2^72^. We then used paftools.js^72^ for variant calling, and vep^73^ for variant effect prediction.

### PGGB Based Graph Construction

While the HPRC’s existing pangenome graphs provide a valuable resource, we discovered that they did not provide the best reference system for genotyping copy number variation. Our validation of the genotyping approach revealed that we would experience high genotyping error when gene copies (e.g. all copies of *AMY1*, or all copies of *AMY2B*) were not fully “collapsed” into a single region in the graph. We thus elected to rebuild the graph locally to improve genotyping accuracy for complex structural variants. This achieves substantially improved results by allowing multiple mappings of each haplotype against others, which leads to a graph in which multi-copy genes are collapsed into single regions of the graph. This collapsed representation is important for graph-based genotyping. Additionally, we incorporated additional samples, some of which were reassembled by us, that were not part of the HPRC’s original dataset to have a more comprehensive representation of variability in the amylase locus, which required rebuilding the pangenome graph model at the amylase locus.

A PGGB graph was constructed from 94 haplotypes spanning the amylase locus using PGGB v0.5.4 (commit 736c50d8e32455cc25db19d119141903f2613a63)^26^ with the following parameters: ‘-n 94‘ (the number of haplotypes in the graph to be built) and ‘-c 2‘ (the number of mappings for each sequence segment). The latter parameter allowed us to build a graph that correctly represents the high copy number variation in such a locus. We used ODGI v0.8.3 (commit de70fcdacb3fc06fd1d8c8d43c057a47fac0310b)^74^ to produce a Jaccard distance-based (i.e. 1-Jaccard similarity coefficient) dissimilarity matrix of paths in our variation graph (‘odgi similarity -d‘). These pre-computed distances were used to construct a tree of relationships between haplotype structures using neighbor joining.

### Haplotype Deconvolution Approach

We implemented a pipeline based on the workflow language Snakemake (v7.32.3) to parallelize *haplotype deconvolution* (i.e., assign to a short-read sequenced individual the haplotype pair in a pangenome that best represents its genotype at a given *locus*) in thousands of samples.

Given a region-specific PGGB graph (gfa, see **PGGB Based Graph Construction**), a list of short-read alignments (BAM/CRAM), a reference build (fasta) and a corresponding region of interest (chr:start-end; based on the alignment of the BAM/CRAM), our pipeline runs as follows:

1. extract the haplotypes from the initial pangenome using ODGI (v0.8.3, `‘odgi paths -f‘`)
2. for each short-read sample, extract all the reads spanning the region of interest using SAMTOOLS (v1.18, ‘`samtools fasta`‘)^75^
3. map the extracted reads back to the haplotypes with BWA (v0.7.17, ‘`bwa mem`‘)^58^. To map ancient samples, we used instead ‘bwa aln‘ with parameters suggested in Oliva A et al., 2021^76^: ‘`bwa aln -l 1024 -n 0.01 -o 2`‘
4. compute a node depth matrix for all the haplotypes in the pangenome: every time a certain haplotype in the pangenome loops over a node, the path depth for that haplotype over that node increases by one. This is done using a combination of commands in ODGI (‘`odgi chop -c 32`‘ and ‘`odgi paths -H`‘)
5. compute a node depth vector for each short-read sample: short-read alignments are mapped to the pangenome using GAFPACK (https://github.com/ekg/gafpack, commit ad31875) and their coverage over nodes computed using GFAINJECT (https://github.com/ekg/gfainject, commit f5feb7b)
6. compare each short-read vector (see .5) with each possible pair of haplotype vectors (see .4) by means of cosine similarity using (https://github.com/davidebolo1993/cosigt, commit e247261) (which measures the similarity between two vectors as their dot product divided by the product of their lengths). The haplotype pair having the highest similarity with the short-read vector is used to describe the genotype of the sample.
7. The final genotypes were assigned as the corresponding consensus haplotypes of highest similarity pair haplotypes.

Our pipeline is publicly available on GitHub (https://github.com/raveancic/graph_genotyper) and is archived in zenodo https://zenodo.org/doi/10.5281/zenodo.10843493.

We assessed the accuracy of the haplotype deconvolution approach in several different ways. First we assessed 35 individuals (70 haplotypes) for which both short-read sequencing data and long-read diploid assemblies were available. In 100% of cases (70/70 haplotypes) we accurately distinguished the correct haplotypes present in an individual from short read sequencing data. We further assessed how missing haplotypes in the pangenome graph might assess the accuracy of our approach by performing a “leave-one-out,” “jackknifing,” analysis. In this approach, for each of the 35 long-read individuals individuals we rebuilt the variation graph with a single haplotype excluded and tested our ability to identify the correct consensus haplotype from the remaining haplotypes. The true positive rate was ∼93% in this case. Second, we compared our haplotype deconvolutions to haplotypes determined by inheritance patterns in 44 families in a previous study (Usher *et al* 2015, Table S3)^15^. We note that this study hypothesized the existence of an H4A4B4 haplotype without having observed it directly. In our study we also find no direct evidence of the H4A4B4 haplotype. Furthermore, we find that inheritance patterns are equally well explained by other directly observed haplotypes and thus exclude these predictions from our comparisons (2 individuals excluded). We identified the exact same pair of haplotypes in 95% of individuals (125/131 individuals) and in 97% of individuals (288/298 individuals) the haplotype pair we identify is among the potential consistent haplotype pairs identified from inheritance. Third, we compared inheritance patterns in 602 diverse short-read sequenced trios from 1000 genomes populations^19^. For each family we randomly selected one parent and assessed if either of the two offspring haplotypes were present in this randomly selected parent. Across all families, this proportion, *p*, represents an estimate of the proportion of genotype calls that are accurate in both the offspring and that parent, thus the single sample accuracy can be estimated as the square root of *p.* From these analyses we identified 533/602 parent-offspring genotype calls that are correct, corresponding to an estimated accuracy of 94%. Fourth, we compared our previously estimated reference genome read-depth-based copy number genotypes to those predicted from *haplotype deconvolutions* across 4,292 diverse individuals. These genotypes exhibited 95-99% concordance across different amylase genes (95%, 97%, and 99% for *AMY1, AMY2A*, and *AMY2B* respectively). Cases in which the two estimates differed were generally high-copy genotypes for which representative haplotype assemblies have not yet been observed and integrated into the graph (**Fig S20**). Overall we thus estimate the *haplotype deconvolution* approach to be ∼95% accurate for modern samples, and thus choose not to propagate the remaining 5% uncertainty into downstream analyses.

To determine the impact of coverage and technical artifacts common in ancient DNA we performed simulations. We selected 40 individuals having both haplotypes represented in the AMY graph and, for those, we simulated short reads mirroring error profiles in modern and ancient genomes across different coverage levels. More specifically, we simulated paired-end short reads for the modern samples with wgsim (https://github.com/lh3/wgsim) (commit a12da33, `‘wgsim −1 150 −2 150`‘) and single-end short reads for the ancient samples with NGSNGS^77^ (commit 559d552, `‘ngsngs -ne -lf Size_dist_sampling.txt -seq SE -m b7,0.024,0.36,0.68,0.0097 -q1 AccFreqL150R1.txt`‘ following author’s suggestions in https://github.com/RAHenriksen/NGSNGS). Synthetic reads were then aligned against the GRCh38 build of the human reference genome using bwa-mem2^78^ (commit 7f3a4db). For samples modeling modern individuals, we generated 5X to 30X coverage data while for those modeling ancient genomes we aimed for lower coverage (1X to 10X) to better approximate true-to-life data. We ran our haplotype deconvolution pipeline independently for modern and ancient simulated samples, as well as varying coverage levels. Out of 480 tests, only 9 (approximately 1%) yielded incorrect predictions, exclusively in ancient simulated sequences with coverage ranging from 1X to 4X. Cosine similarity scores for ancient simulated sequences ranged from 0.789 to 0.977 (median=0.950), while scores for modern simulated sequences ranged from 0.917 to 0.992 (median=0.981) (**Fig S23**). We therefore conclude that the haplotype deconvolution method is highly accurate for ancient samples as well. Out of an abundance of caution, we further imposed a conservative quality score threshold of 0.75 to ancient samples, resulting in 288 ancient samples with high-confidence haplotype assignment out of a total of 533 (**Figs S24-S25**). We note that the haplotype deconvolutions in ancient samples are likely more accurate than read depth genotypes which tend to be biased towards higher copy number.

### LD estimation

To investigate pairwise linkage disequilibrium (LD) across the SVR region at a global scale, we first merged our copy number estimates with the joint SNP call set from HGDP and 1kGP^57^, resulting in a variant call set of 3,395 diverse individuals with both diploid copy number genotypes and phased SNP calls. Briefly, we used bcftools-v1.9^75^ to filter HGDP and 1kGP variant data for designated genomic regions on chromosome 1, including the amylase structurally variable region (SVR) and flanking regions defined as bundle 0 and bundle 1 (distal and proximal respectively) using the GRCh38 reference coordinate system (--region chr1:103456163-103863980 in GRCh38). The resulting output was saved in variant call format (vcf), keeping only bi-allelic SNPs (-m2 -M2 -v snps), and additionally filtered with vcftools-v.0.1.16^79^ with –keep and –recode options for lists of individuals grouped by continental region in which we were able to estimate diploid copy numbers. Population-specific vcf files were further filtered for a minor allele frequency filter threshold of 5% (--minmaf 0.05) and used to generate a numeric genotype matrix with the physical positions of SNPs for LD calculation (R^2^ statistic) and plotting with the LDheatmap^80^ function in R-v4.2.2.

To further dissect the unique evolutionary history of the amylase locus, we compared regions with high R^2^ across the SVR with LD estimates for pairs of SNPs across regions of similar size in chromosome 1. We specifically focused on pairs of SNPs spanning bundle 0 (chr1:103456163-103561526 in GRCh38) and the first 66-kbp of bundle 1, hereafter labeled as bundle 1a (chr1:103760698-103826698 in GRCh38), as revealed by the LD heatmap. Then we computed the R^2^ values for any pair of SNPs in chromosome 1 for each superpopulation within a minimum of 190 kb distance (i.e. the equivalent distance from bundle 0 end to bundle 1a start using the GRCh38 reference coordinate system) and maximum 370 kb distance (i.e.the equivalent distance from bundle 0 start to bundle 1a end using the GRCh38 reference coordinate system). To calculate pairwise LD across the human chromosome 1 for different populations we ran plink-v1.90b6.21^81^ with options -r2 –ld-window 999999 –ld-window-kb 1000 –ld-window-r2 0 – make-bed –maf 0.05, using as input population-specific vcf files for a set of biallelic SNPs of 3,395 individuals from HGDP and 1kGP. Since the resulting plink outputs only provide R^2^ estimates for each pair of SNPs and respective SNP positions, we additionally calculated the physical distances between pairs of SNPs as the absolute difference between the base pair position of the second (BP_B) and first (BP_A) SNP. We then filtered out distances smaller than 190 kb and greater than 370 kb, and annotated the genomic region for each R^2^ value based on whether both SNPs fall across the SVR region or elsewhere in chromosome 1. The distance between SNP pairs was also binned into intervals of 20,000 bp and each interval’s midpoint was used for assessing LD decay over genomic distances. The resulting dataset was imported in R to compute summary statistics comparing LD across each major continental region, or superpopulations, and we used ggplot2 to visualize the results.

### Coalescent tree, ancestral state reconstruction, and PCA

To construct the coalescent tree, we first extracted bundle 0 and bundle 1a sequences from all 94 haplotypes (i.e. distal and proximal unique regions flanking the amylase SVR) that went through principal bundle decomposition. Based on their coordinates on the human reference genome (GRCh38), we used samtools-v1.17^82^ to extract these sequences from three Neanderthal and one Denisovan genomes that are aligned to GRCh38. We used kalign-v3.3.5^83^ to perform multiple sequence alignment on bundle 0 and bundle 1a sequences. We used iqtree-v2.2.2.3^84^ to construct a maximum likelihood tree with Neanderthal and Denisova sequences as the outgroup, using an estimated 650 kyr human-Neanderthal split time for time calibration ^29^. We used ggtree-v3.6.2^85^ in R-v4.2.1 to visualize the tree, and annotated each tip with its structural haplotype and amylase gene copy numbers. We used cafe-v5.0.0^86^ to infer the ancestral copy numbers of each of the three amylase genes along the time-calibrated coalescent tree (excluding the outgroups) and to estimate their duplication/deletion rates. The timing of each duplication/deletion event was estimated based on the beginning and end of the branch along which the amylase gene copy number has changed. We used ggtree and ggplot-v3.4.2 in R to visualize these results, and used Adobe Illustrator to create illustrations for several of the most notable duplication/deletion events^87^.

Next, we performed a principal component analysis (PCA) combining 94 HPRC haplotype sequences with variant calls for 3,395 individuals from HGDP and 1kGP. We first aligned all 94 bundle 0 and 94 bundle 1a haplotype sequences to the human reference genome (GRCh38) using minimap2-v2.26^72^, and called SNPs from haplotypes using paftools.js. Each haplotype sequence appears as a pseudo-diploid in the resulting vcf file (i.e. when the genotype is different from the reference, it is coded as being homozygous for the non-reference allele). These haplotype-specific vcf files were merged together and filtered for biallelic SNPs (-m2 -M2 -v snps) with bcftools, resulting in a pseudo-diploid vcf file from 94 haplotype sequences for each bundle. These were then merged with the respective bundle 0 and bundle 1a vcf files from HGDP and 1kGP, also filtered for biallelic SNPs, using bcftools. Finally, we ran plink with a minor allele frequency of 5% (--maf 0.05) to obtain eigenvalues and eigenvectors for PCA and used ggplot-v3.4.2 to visualize the results. These analyses were conducted with bundle 0 and bundle 1a separately, with highly concordant results (**Figs S6-S7**). Analyses focused on bundle 0 are reported in the main text (**Fig 3**) whereas bundle 1a results are shown as extended data (**Extended Data Figure 3**).

### Signatures of recent positive selection in modern human populations

To investigate very recent or ongoing positive selection at the amylase locus in modern humans, we first looked for significant signatures of reduced genetic diversity across the non-duplicated regions adjacent to the SVR compared to chromosome 1 in different populations worldwide. This stems from the assumption that, given low SNP density across the SVR, the high levels of LD found between pairs of SNPs spanning bundle 0 and bundle 1a indicate that SNPs in bundle 0 or bundle 1 can be used as proxies for the selective history of the linked complex structures of the SVR. We calculated nucleotide diversity (π) on sliding windows of 20,000 bp spanning GRCh38 chromosome 1 with vcftools using as input population-specific vcf files from HGDP and 1kGP filtered for a set of biallelic SNPs. Each window was annotated for the genomic region, namely bundle 0, SVR and bundle 1a. All windows comprising the SVR region were removed from the resulting output due to low SNP density. We then used ggplot2 in R to calculate and visualize summary statistics comparing nucleotide diversity for windows located windows harboring the flanking regions to amylase genes (i.e. bundle 0 and 1a) with nucleotide diversity for windows spanning the rest of chromosome 1 for each major continental region or super population.

To identify either soft and hard selective sweeps at the flanking regions of the SVR, we computed several different extended haplotype homozygosity-based statistics and statistics based on distortions of the haplotype frequency spectrum (**Table S5**). Vcf files from HGDP and 1kGP chromosome 1-22 GRCh38 were filtered for biallelic SNPs and minor allele frequency of 0.05 for target populations with over 10 individuals to calculate iHS ^31^, nSL ^32^, XP-nSL ^35^ as implemented in *selscan* ^88^ (see **Table S5** for a description of populations and selection statistics). CEU and YRI populations were also included to confirm the ability of the tests to consistently identify the LCT hard sweep in CEU and in relation to the amylase locus (**Table S5**). Scores for these statistics were normalized using the genome-wide empirical background with selscan’s co-package *norm.* This was also used to compute the fraction of the standardized absolute values > 2 for each statistic in non-overlapping 100kb windows genome-wide^31^. For XP-nSL statistics, modern rainforest hunter-gatherers in Africa and the pastoralists Yakut were used as reference populations, so that positive scores correspond to possible sweeps in the populations with traditionally agricultural diets. We additionally used *lassip*^34^ to compute H12 and H2/H1 statistics^33^ and saltiLASSI Λ^34^ on sliding windows of 201 SNPs with intervals of 100 SNPs. SNP positions within the SVR region were removed from the resulting outputs due to low SNP density. We then compared the average and distribution of all selection statistics across individual SNPs or windows located within bundle 0 and bundle 1a (labeled as ‘AMY region’) and located within chr2:135Mb-138Mb (labeled as ‘LCT region’) with that of the rest of the genome using geom_stats() and geom_density() functions in ggplot2 (**Fig S8-S19, Table S5**). We also used an outlier approach, and focused on the top 0.05% of the test statistic across all windows genome-wide for modern populations of known subsistence, and considered estimates above this threshold to be strong signals of selection^31^.

### Inference of recent positive selection in West Eurasian populations using ancient genomes

To determine if changes in the frequency of different structural haplotypes over the last 12,000 years were consistent with positive selection, we first grouped amylase structural haplotypes (n=11) into those with the ancestral number of amylase gene copies (three total), or with amylase gene duplications (five or more copies). We used three complementary approaches to infer the selection coefficient associated with duplication-containing haplotypes. First, we used ApproxWF^38^ to perform Bayesian inference of the selection coefficient from binned allele frequency trajectories. We ran ApproxWF for 1010000 MCMC steps with parameters N=10000, h=0.5, and pi=1. We assumed a generation time of 30 years to convert the age of ancient samples from years to generations. The first 10,000 steps of the MCMC process were discarded in all analyses. Next, we used bmws^39^ to estimate the allele frequency trajectory and time-varying selection from genotype data with parameters -d diploid -l 4.5 -g 30 -n 10000 -t. We further ran 1000 bootstrap replicates to obtain 95% credible intervals around our estimates. Lastly, we used an ABC approach adapted and modified from^40^ to explicitly account for the demographic processes underlying the allele frequency changes. We performed extensive forward-in-time simulations using SLiM^89^ based on a well-established demographic model for West Eurasians^45^ that includes major population split and admixture events as well as population growth (**Table S11**). We allowed three model parameters to vary across simulations: selection coefficient (*s*), the time of selection onset (*t*, in kyr BP), and the initial allele frequency in the ancestral population (*f*). Selection is only applied to known agricultural populations (i.e., Early farmers, Neolithic farmers, and Bronze age to present day Europeans), and its strength is assumed to be constant over time. These parameter values were set in evenly spaced intervals (i.e., 21 values of *s* ∈ [-0.01, 0.04], 21 values of *t* ∈ [3, 15], 31 values of *f* ∈ [0.05, 0.8]), and 1,000 replicate simulations were run for each unique parameter combination. This resulted in 13,671,000 simulations in total. For each simulation, we calculated the difference between the observed and the expected binned allele frequency trajectories, accounting for uneven sampling in time and genetic ancestry. We then selected the top 0.1% of simulations (i.e. 13,671 simulations) that best ressemble the observed data to approximate the posterior distribution of model parameters. We also examined the allele frequency changes (i.e. the difference between allele frequencies in the first and last time bin) across all neutral simulations with s=0 and compared them to the observed allele frequency change in the data (**Fig S29**).

## Data Availability

All data used in this project are publically available and described in the Datasets section of the methods. Copy number genotypes, structural haplotypes, haplotype deconvolutions, and pangenome graphs can be found in Supplementary Tables and the zenodo archived github repository.

## Code Availability

All code can be found deposited in the following GitHub repository https://github.com/sudmantlab/amylase_diversity_project and is archived in zenodo 10.5281/zenodo.10995434.

## Acknowledgements

We would like to thank Morten E. Allentoft, Rasmus Nielsen, Evan K. Irving-Pease, Martin Sikora, Joshua Schraiber, Vince Buffalo, and Eske Willersvlev, for helpful discussion and assistance in accessing ancient datasets. Institute of General Medical Sciences [grant: R35GM142916] to PHS. Vallee Scholars Award to PHS. Ancient DNA sequencing was supported by grants from the Lundbeck Foundation (R302-2018-2155 and R155-2013-16338).

## Contributions

Conceived the experimental design: PHS Processed and analyzed the data: PHS, DB, AH, RNL, AR, JR, AG, JC, EG Wrote and edited the manuscript: PHS, DB, AH, RNL, AR, JR, AG, JC, EG Supervised research: PHS, EG, NS

---

**Extended Data Figure 1 -.**
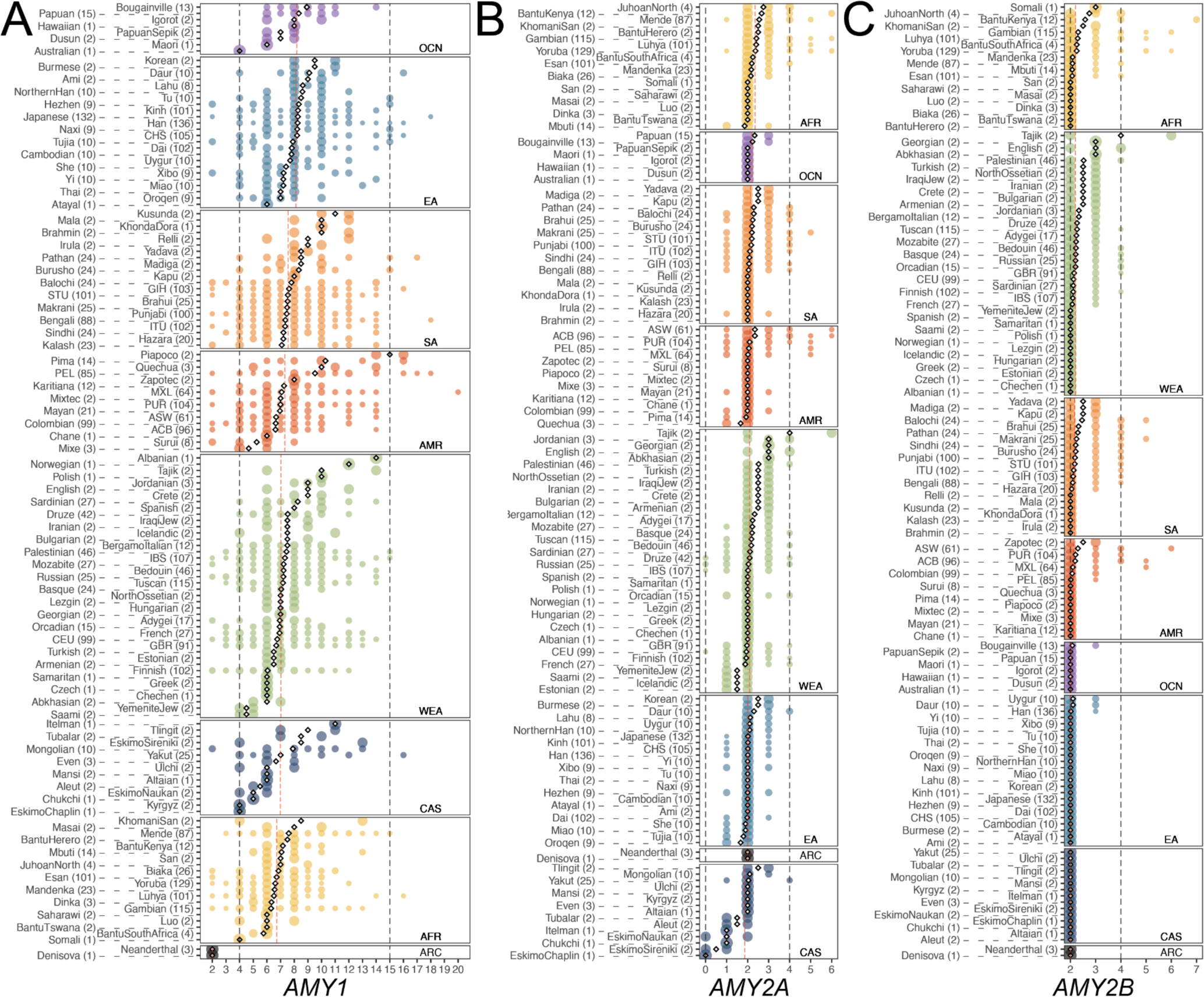
Worldwide amylase subpopulation copy number diversity. **A-C)** Copy number distributions of *AMY1, AMY2A*, and *AMY2B* in 147 modern human populations and four archaic hominids. The size of each point is proportional to the proportion of individuals in the population with that genotype. Diamonds indicate the subpopulation mean, red dashed lines indicate the continental population mean, grey dashed lines indicate minimum and maximum subpopulation means.

**Extended Data Figure 2 -.**
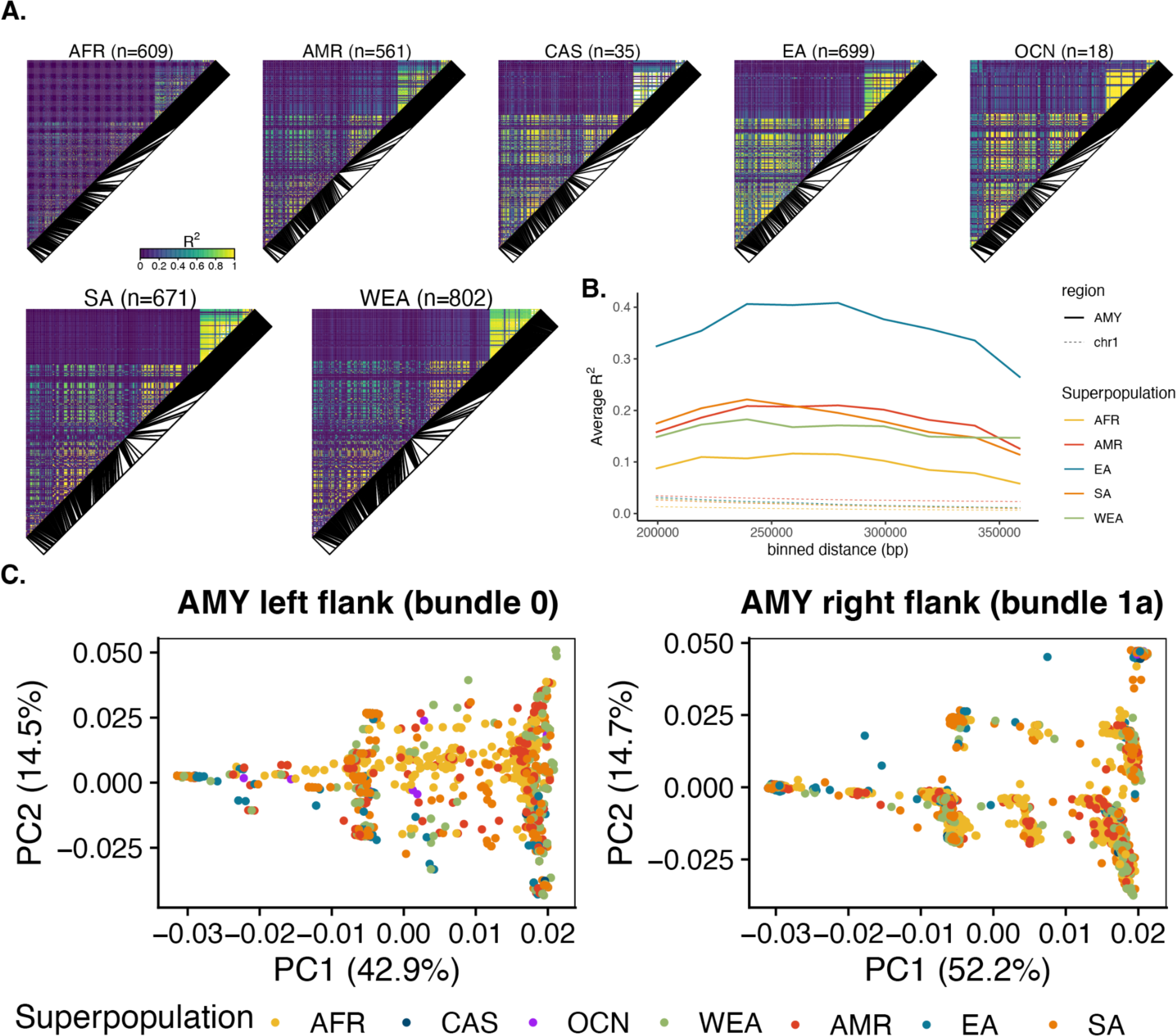
LD in different populations worldwide including 3,395 diverse diploid human genomes. **A)** Heat maps of linkage disequilibrium (LD) for SNPs across a ∼406 kb region spanning unique sequences on either side of the structurally variable region of amylase (SVR) in different populations from seven continental regions (Africa - AFR, America - AMR, Central Asia - CAS, East Asia - EA, Oceania - OCN, South Asia - SA and Western Eurasia - WEA). **B)** LD decay over genomic distances for groups with more than 100 samples, measured as the average R2 between SNP pairs on either side of the SVR (i.e. 190 kb - 370 kb apart) binned into intervals of 20,000 bp, compared to identically spaced SNPs in chromosome 1. **C)** PCAs for non-duplicated regions adjacent to the SVR according to different continental regions using the distal (bundle 0) and proximal (bundle 1a) regions (see also **Figure S7**).

**Extended Data Figure 3 -.**
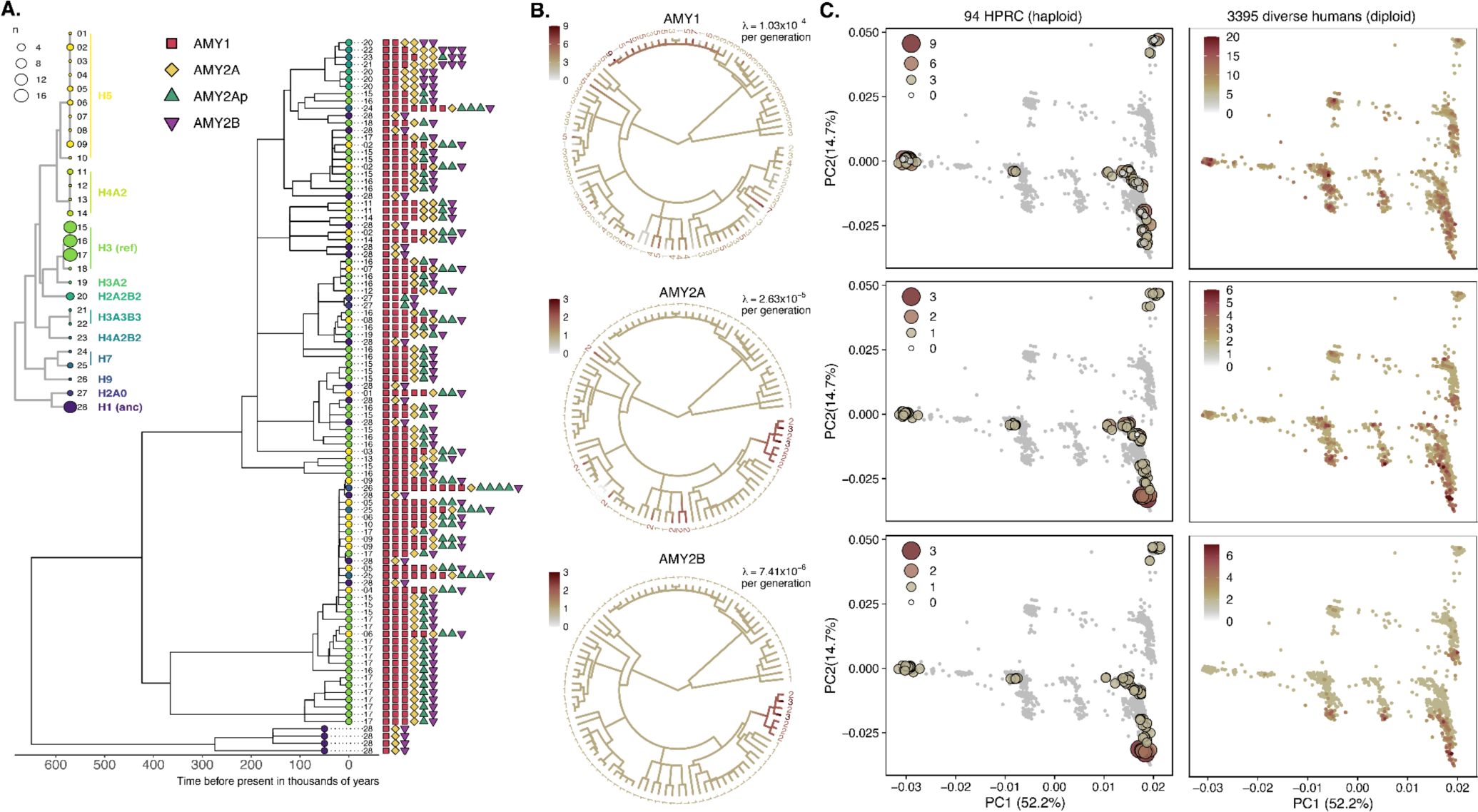
Reconstruction of the evolutionary history of amylase structural haplotypes using proximal unique sequence. **A)** A time-calibrated coalescent tree from the proximal non-duplicated region flanking the SVR (rightmost gray arrow in A until the recombination hotspot) across 94 assembled haplotypes (tree from the distal region in **Fig. 3**). The number next to each tip corresponds to the structural haplotype that the sequence is physically linked to and the color of the circle at each tip corresponds to its consensus haplotype structure (see inset structure tree). The copy numbers of each amylase gene and pseudogene are also shown next to the tips of the tree. **B)** Ancestral state reconstruction and mutation rate estimates for amylase gene copy number (archaic outgroups excluded). Branch color corresponds to copy number. **C)** A PCA from 94 haplotype assemblies and 3,395 diverse diploid human genomes from the proximal non-duplicated region flanking the SVR (PCA from the distal region in **Fig. 3**).

